# Intestinal transkingdom analysis on the impact of antibiotic perturbation in health and critical illness

**DOI:** 10.1101/2020.06.25.171553

**Authors:** Bastiaan W. Haak, Ricard Argelaguet, Cormac M. Kinsella, Robert F.J. Kullberg, Jacqueline M. Lankelma, Theodorus B.M. Hakvoort, Floor Hugenholtz, Sarantos Kostidis, Martin Giera, Wouter J. de Jonge, Marcus J. Schultz, Tom van Gool, Tom van der Poll, Willem M. de Vos, Lia van den Hoek, W. Joost Wiersinga

**Affiliations:** Center for Experimental and Molecular Medicine, Amsterdam UMC, Location AMC, Amsterdam Infection & Immunity Institute, Amsterdam, The Netherlands; European Molecular Biology Laboratory, European Bioinformatics Institute, Hinxton, Cambridge, United Kingdom; Laboratory of Experimental Virology, Department of Medical Microbiology, Amsterdam UMC, Location AMC, Amsterdam, the Netherlands; Center for Proteomics and Metabolomics, Leiden University Medical Center, Leiden, The Netherlands; Tytgat Institute for Liver and Intestinal Research, Amsterdam UMC, Location AMC, Amsterdam, The Netherlands; Department of Intensive Care, Amsterdam UMC, Location AMC, Amsterdam, The Netherlands; Department of Parasitology, Amsterdam UMC, Location AMC, Amsterdam, The Netherlands; Laboratory of Microbiology, Wageningen University, Wageningen, The Netherlands; Research Programs Unit Immunobiology, Department of Bacteriology and Immunology, Helsinki University, Helsinski, Finland; Department of Medicine, Division of Infectious Diseases, Amsterdam UMC, Location AMC, Amsterdam, The Netherlands

## Abstract

Bacterial microbiota play a critical role in mediating local and systemic immunity, and shifts in these microbial communities have been linked to impaired outcomes in critical illness. Emerging data indicate that other intestinal organisms, including bacteriophages, viruses of eukaryotes, fungi, and protozoa, are closely interlinked with the bacterial microbiota and their host, yet their collective role during antibiotic perturbation and critical illness remains to be elucidated. Here, multi-omics factor analysis (MOFA), a novel computational strategy to systematically integrate viral, fungal and bacterial sequence data, we describe the functional impact of exposure to broad-spectrum antibiotics in healthy volunteers and critically ill patients. We observe that a loss of the anaerobic intestinal environment is directly correlated with an overgrowth of aerobic pathobionts and their corresponding bacteriophages, as well as an absolute enrichment of opportunistic yeasts capable of causing invasive disease. These findings further illustrate the complexity of transkingdom interactions within the intestinal environment, and show that modulation of the bacterial component of the microbiome has implications extending beyond this kingdom alone.

## Introduction

In recent years, widespread efforts have been dedicated on elucidating the immunomodulatory impact of intestinal microorganisms in health and disease (Belkaid & Hand, 2014; Honda & Littman, 2012). Animal studies have shown that broad-spectrum antibiotic modulation of the intestinal microbiota enhances susceptibility to enteric and systemic infections (Schuijt *et al*, 2015; Clarke *et al*, 2010; Buffie & Pamer, 2013). In line with these preclinical findings, our group and others have observed that exposure to broad-spectrum antimicrobial therapy profoundly distorts the composition of the intestinal microbes of critically ill patients in the Intensive Care Unit (ICU) (Lankelma *et al*, 2017b; McDonald *et al*, 2016; Zaborin *et al*, 2016). These disruptions within the intestinal environment enable the rapid expansion of opportunistic pathobionts and nosocomial infections, including infections with vancomycin-resistant enterococci as well as invasive disease by antibiotic-resistant Enterobacteriaceae (Haak & Wiersinga, 2017; Taur *et al,* 2012; Agudelo-Ochoa *et al*, 2020).

Traditionally, viruses were considered solely pathogens; however, growing evidence suggests a more dynamic relationship between the virome and the host, mediated through direct interactions with the bacterial microbiome(Shkoporov & Hill, 2019; Norman *et al*, 2014; Pfeiffer & Virgin, 2016; Neil & Cadwell, 2018). Viruses influence immune development and shape tissue architecture (Kuss *et al*, 2011; De Sordi *et al*, 2019), and changes in the composition of viral communities have been associated with disease severity in inflammatory bowel disease (IBD), acquired immune deficiency syndrome (AIDS), and the development of Graft versus Host Disease (GvHD) (Shkoporov & Hill, 2019; Legoff *et al*, 2017; Zuo *et al*, 2019). Similarly, intestinal fungi have recently been acknowledged as a small but potentially important part of the intestinal ecosystem and have been shown to play a potentially immunomodulatory role in the development of colorectal cancer, IBD, and irritable bowel syndrome (IBS) (Sokol *et al*, 2017; Botschuijver *et al,* 2017; Sovran *et al,* 2018; Richard & Sokol, 2019). In addition, a recent study reported that a reduction of anaerobic bacteria during the course of allogeneic hematopoietic stem cell transplantation directly facilitates the intestinal overgrowth of specific *Candida* species, ultimately culminating in invasive fungal disease (Zhai *et al*, 2020).

While these findings provide clues that specific cross-kingdom interactions potentially contribute to or exacerbate disease, a large knowledge gap remains on the composition, interactions and functions of fungi and viruses following exposure to broad-spectrum antibiotics, both in healthy volunteers and in patients with critical illness. Hence, there is an increasing need for unsupervised integrative computational frameworks that can robustly and systematically identify underlying patterns of variation across these communities in health and disease (Shkoporov & Hill, 2019; Richard & Sokol, 2019).

## Results and Discussion

To examine the extent of these transkingdom interactions during critical illness, we collected faecal samples from 33 patients (mean age 64 years; 48% male; **Table EV1**) admitted to the Intensive Care Unit (ICU) of the Amsterdam University Medical Centres, location Academic Medical Centre, Amsterdam, the Netherlands. Of these patients, 24 were admitted with sepsis while nine patients had a non-infectious diagnosis (non-septic ICU). All ICU patients were treated with between one and nine different classes of antimicrobial agents (**Fig. EV1**). Thirteen healthy non-smoking volunteers were evaluated as controls. Six healthy subjects received oral broad-spectrum antibiotics (ciprofloxacin 500 mg q12h, vancomycin 500 mg q8h and metronidazole 500 mg q8h) for seven days, whereas seven subjects did not receive antibiotics. Subjects were asked to collect faecal samples before antibiotic treatment and one day after completing the course of antibiotics.

We performed sequencing of the V3-V4 region of the bacterial 16S ribosomal RNA (rRNA) gene and the fungal Intergenic Transcribed Spacer (ITS)1 rRNA gene, seeking to examine community compositions by characterizing fungal and bacterial sequences into exact amplicon-sequencing variants (ASVs) (Callahan *et al*, 2016). We simultaneously performed virus discovery next-generation sequencing (VIDISCA-NGS) (van der Hoek *et al*, 2012) using a validated virome-enriched library preparation (Edridge *et al*, 2019; Kinsella *et al*, 2019). Finally, we measured the presence or absence of intestinal gut protozoa using targeted polymerase chain reaction (PCR).

The bacterial microbiome of ICU patients and volunteers exposed to antibiotics included in this study has been described previously by our group (Lankelma *et al*, 2017b; Haak *et al*, 2019) (**Fig. 1a**). Bacterial alpha diversity and richness dropped significantly in ICU patients and healthy subjects exposed to antibiotics, with the latter most significantly impacted in both metrics (**Fig. 1b**). In line with earlier observations (Hallen-adams & Suhr, 2017; Suhr & Hallen-Adams, 2015; Nash *et al*, 2017), fungal communities were dominated by *Candida* and *Saccharomyces*, while *Malassezia* and *Aspergillus* were also frequently observed. Overall, fungal diversity metrics were comparable between critically ill patients and healthy controls not exposed to antibiotics, while significant drops in diversity were observed in healthy subjects after exposure to antibiotics. Viral communities were largely dominated by environmental single stranded (ss)RNA viruses and bacteriophages of the order Caudovirales. Strikingly, around 50% of the abundance of the virome consisted of cross-assembly (crAss) phages, which have recently been connected to *Bacteroides* spp (Dutilh *et al*, 2014; Shkoporov *et al*, 2018). No differences in viral alpha diversity were observed, yet both septic ICU patients and antibiotic perturbed volunteers displayed higher viral richness. We observed short-term temporal stability of all three kingdoms in healthy subjects not receiving antibiotics (Shkoporov *et al*, 2019) (**Fig. EV2**). In line with recent studies (van Hattem *et al,* 2017, 2019), we observed that a total of 30% of healthy subjects were colonized by the anaerobic gut protozoa *Blastocystis hominis* or *Dientamoeba fragilis*, yet these protozoa were undetectable following antibiotic administration (**Table EV2**).

**Figure 1:**
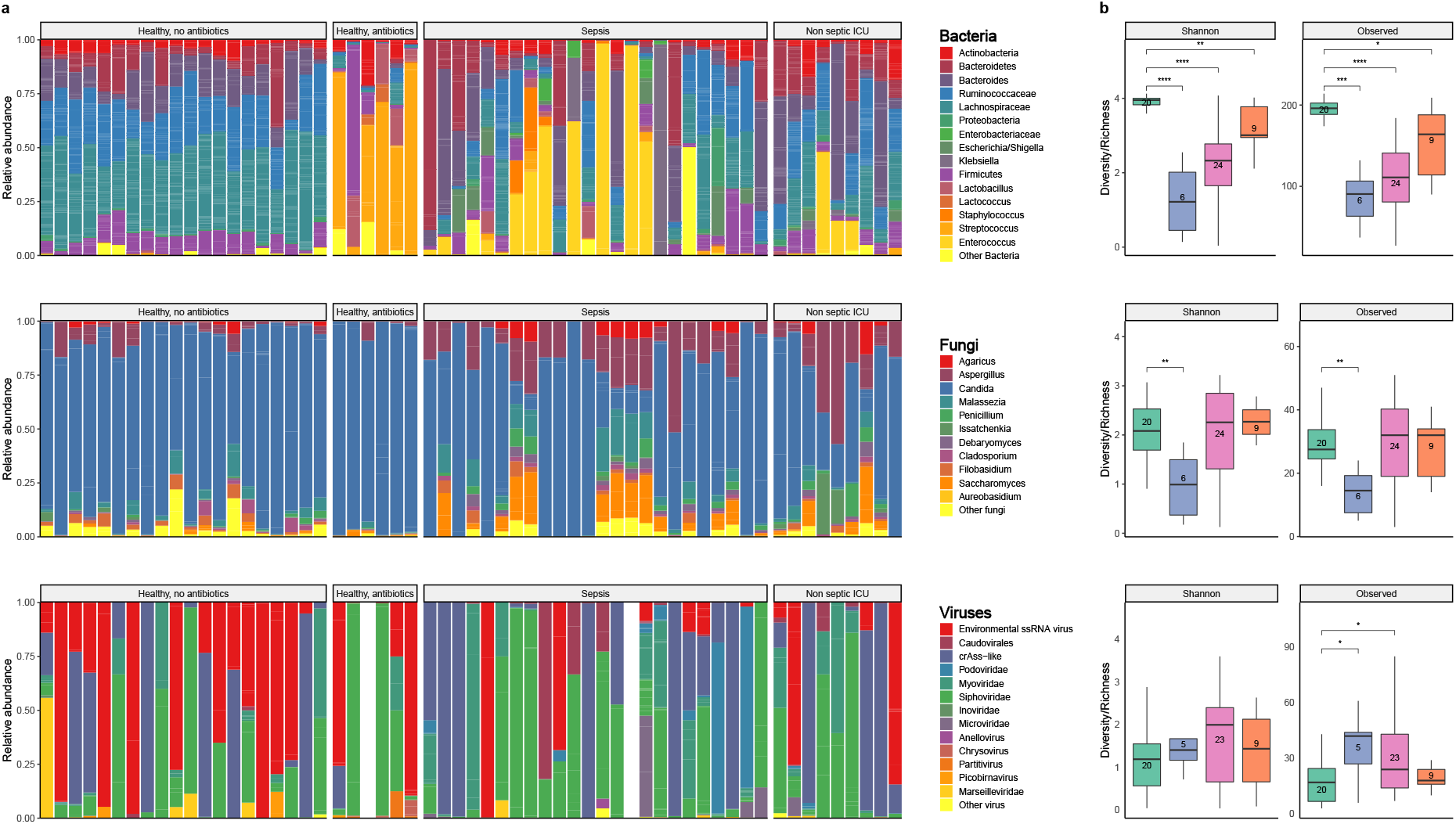
Overview of the composition and diversity of the bacterial, fungal and viral microbiome. (a) Relative proportion of sequence reads at the Genus level assigned to different bacterial and fungal taxa and Order level for viral taxa. Viral metagenomics of two samples did not pass quality control due to high background levels, and were therefore excluded from further analysis. A subset of the bacterial 16S rRNA sequencing data has been previously reported.25,31 (b) Alpha diversity metrics of bacteria (top), fungi (middle) and viruses (bottom), using the Shannon Diversity Index (Shannon) and Observed Taxa richness index (Observed). In the box plots, the central rectangle spans the first quartile to the third quartile (the interquartile range or IQR), the central line inside the rectangle shows the median, and whiskers above and below the box. Given the nonparametric nature of the data, p values were calculated using the Wilcoxon rank sum test.

In order to further understand the patterns of covariation between these intestinal communities during health and critical illness, we employed multi-omics factor analysis (MOFA), a recently developed computational framework for data integration (Argelaguet *et al*, 2018, 2019a). Briefly, MOFA performs unsupervised matrix factorisation simultaneously across multiple data modalities, thereby capturing the global sources of variability via a limited number of inferred factors, effectively yielding a compressed low-dimensional representation of the data. Importantly, the model disentangles the patterns of covariation that are shared across data modalities from the variation that is exclusive to a single data modality (Argelaguet *et al*, 2018) (**Fig. 2a**). This integrative strategy, initially developed for the analysis of single-cell assays (Argelaguet *et al*, 2019b), is especially effective for the analysis of sparse readouts, including microbiome data. As input to the model, we collapsed the inferred bacterial and fungal ASVs and viral reads to their respective Family or Genus level. The number of sequences were subsequently scaled using a centralized-log ratio (Aitchison, 1982), which has shown to be effective in normalizing compositional data (Gloor *et al*, 2017). MOFA identified six Factors with a minimum explained variance of 5% (see Materials & Methods). Altogether, the latent representation explained 39% of the sample heterogeneity in bacteria, 39% for fungi and 19% for viral composition (**Fig. 2b,c**; **Fig. EV3**). Notably, Factor 1 and Factor 3 (sorted by variance explained) captured coordinated variability across all three kingdoms and were capable of completely partitioning transkingdom signatures pertaining to critical illness, antibiotic perturbation and health (**Fig. 2d**). The four remaining Factors identified sample heterogeneity related to low abundant fungal variations (Factor 2; **Fig. EV4**), fluorquinolone/cephalosporin exposure (Factor 4; **Fig. EV5**), as well as bacterial (Factor 5) and viral (Factor 6) signatures pertaining to individual ICU patients.

**Figure 2:**
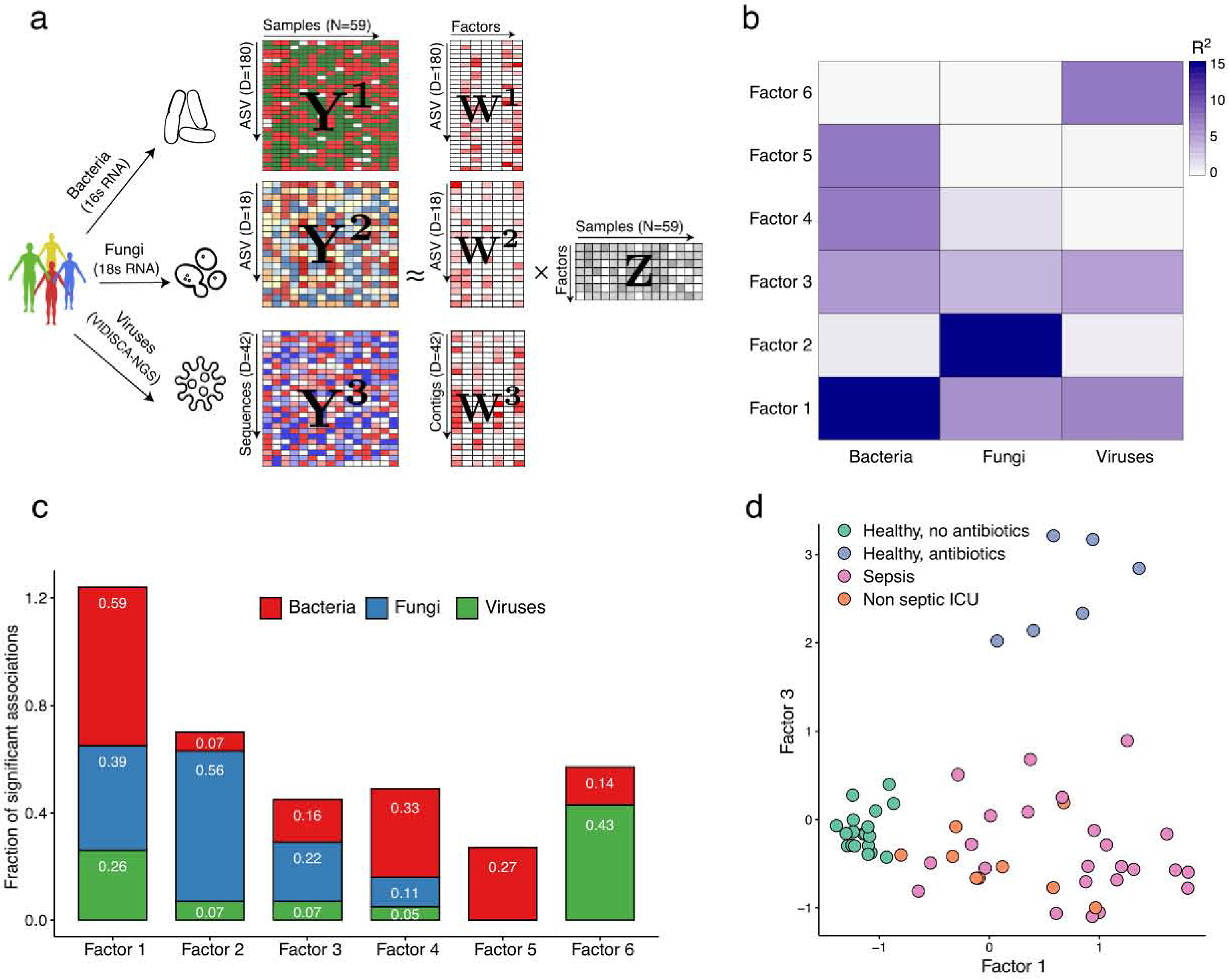
Multi-Omics Factor Analysis (MOFA) delineates the sources of cross-kingdom heterogeneity in the cohort. (a) Model overview: MOFA takes as input the three microbiome quantification matrices. MOFA exploits the covariation patterns between the features within and between microbiome modalities to learn a low-dimensional representation of the data in terms of a small number of latent Factors (Z matrix) and three different weight matrices (W, one per kingdom). By maximising the variance explained under sparsity assumptions,11,12 MOFA provides a principled way to discover the global sources of variability in the data. For each latent Factor (i.e. each source of variation), the weights provide a measure of feature importance for every feature in each Factor, hence enabling the interpretation the variation captured by every factor. (b) Heatmap displays the percentage of variance explained (R2) by each Factor (rows) across the three microbe modalities (columns). Factors 1 and 3 capture coordinated variation across all three microbiome modalities, whereas Factor 2, 4 and 5 are mostly dominated by heterogeneity in Fungi composition. (c) Bar plots show the fraction of significant associations between the features of each microbiome modality and each factor. P-values are obtained using a t-test based on the Pearson’s product moment correlation coefficient. Statistical significance is called at 10% FDR. This plot is useful to interpret whether the variance explained values displayed in (b) are driven by a strong change in a small number of features, or by a moderate effect across a large range of features. (d) Scatter plot of Factor 1 (x-axis) versus Factor 3 (y-axis). Each dot represents a sample, coloured by condition. Factor 1 captures the gradient in microbiome variation associated with antibiotic treatment and critical illness (from negative to positive Factor values), whereas Factor 3 captures the variation associated with antibiotic treatment in healthy patients (positive Factor 3 values) versus critically ill patients (negative Factor 3 values).

Factor 1, the major source of variation, was linked to a transkingdom signature driven by antibiotic perturbation in both health and critical illness, while being consistently absent in healthy subjects without antibiotic exposure (**Fig. 3a,b**). Specifically, bacterial taxa positively associated with this factor were facultative aerobic bacterial pathobionts that have been previously associated with critical illness(Alverdy & Krezalek, 2017; Wischmeyer *et al*, 2016; Haak *etal,* 2017), such as *Staphylococcus, Enterococcus, Klebsiella, Escherichia/Shigella* and *Enterobacter.* Bacterial taxa that were negatively associated with this factor consisted predominantly of genera within the obligatory anaerobic families Lachnospiraceae and Ruminococcaceae, which have been identified as markers of a healthy microbiota and are linked to colonization resistance against bacterial pathobionts(Lee *et al*, 2017; Taur *et al*, 2012). Fungal taxa positively associated with this factor were characterized by yeasts capable of causing invasive disease, such as *Candida, Aspergillus,* and *Debaryomyces* (Miceli *et al*, 2011; Beyda *et al*, 2013; Zhai *et al*, 2020), with a relative absence of the gut constituents *Filobasidium, Malassezia* and *Dipodascus* (Suhr & Hallen-Adams, 2015). The specific cooccurrences of fungal and bacterial taxa observed in Factor 1 are supported by previous studies. For example, members of the Lachnospiraceae family, such as *Blautia* and *Roseburia,* display a direct inhibitory effect on the growth of several *Candida* spp. and *Saccharomyces cerevisiae*, through the production of short-chain fatty acids (SCFAs) and other metabolites (Nguyen *et al*, 2011; García *et al*, 2017; Fan *et al*, 2015). In addition, *in vitro* studies have shown that metabolites produced by *Candida* spp. enhance the growth of *E. coli* and *S. aureus* (Huseyin *et al*, 2017; Kong *et al*, 2017), providing further indications that the intestinal transkingdom signatures identified by MOFA are biologically meaningful.

**Figure 3:**
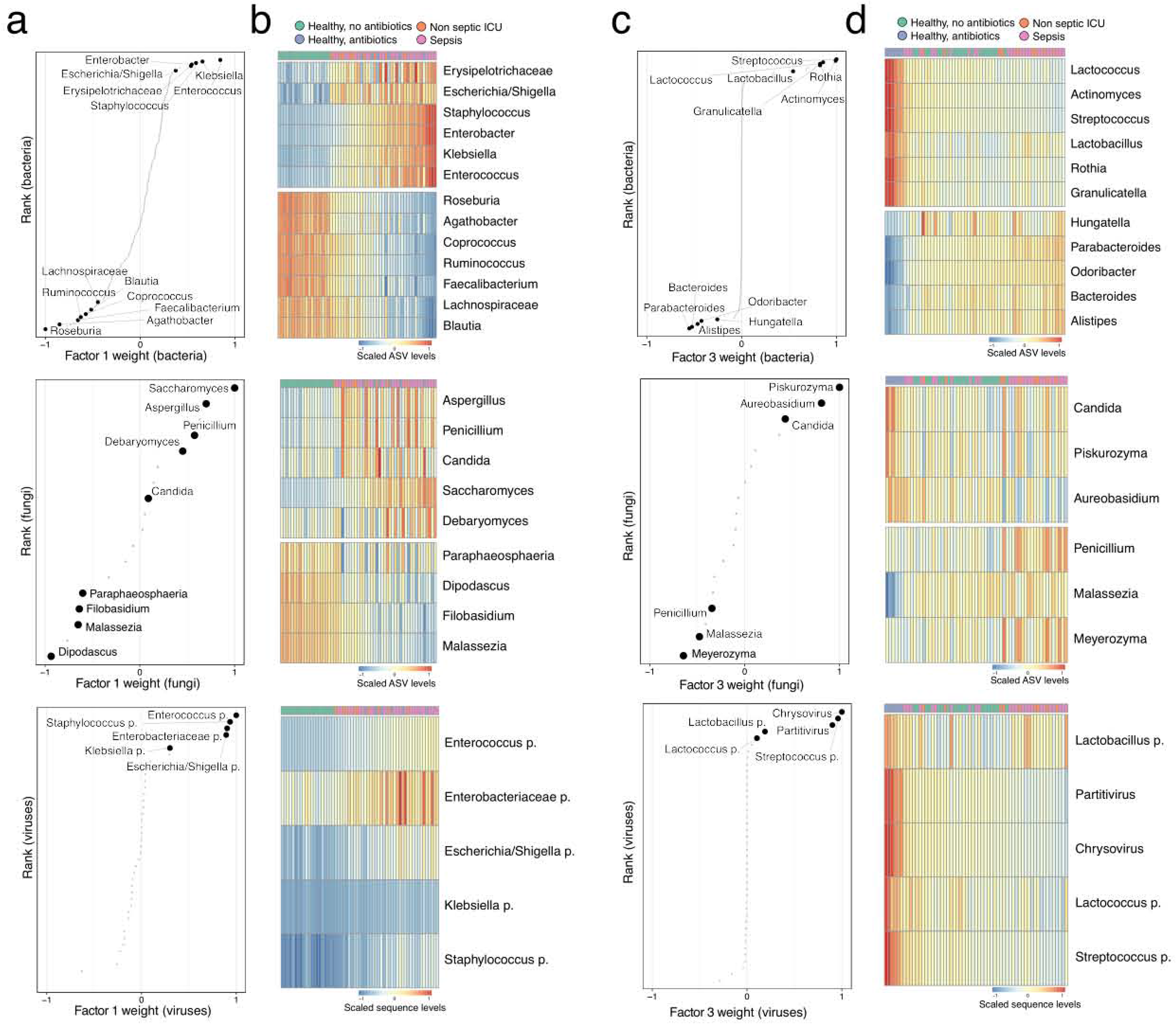
Characterisation of the transkingdom variation captured by Factor 1 and Factor 3. (a) Scatter plots display the distribution of Bacterial (top), Fungi (middle) and Viruses (bottom) weights for Factor 1. A positive value indicates a positive association with Factor 1 values, whereas a negative value indicates a negative association with Factor 1 values (see Fig 2d). The larger the absolute value of the weight, the stronger the association. For ease of visualisation, weights are scaled from −1 to 1. Representative taxa among the top weights are labeled. (b) Heat maps display the reconstructed data (see Methods) based on the MOFA model for the taxa highlighted in (a). Samples are shown in the columns and features in the rows. (c) Scatter plots display the distribution of Bacterial (top), Fungi (middle) and Viruses (bottom) weights for Factor 3. A positive value indicates a positive association with Factor 3 values, whereas a negative value indicates a negative association with Factor 3 values (see Fig 2d). The larger the absolute value of the weight, the stronger the association. For ease of visualisation, weights are scaled from −1 to 1. Representative taxa among the top weights are labeled. (d) Heat maps display the (denoised) data reconstruction (see Methods) based on the MOFA model for the taxa highlighted in (c). Samples are shown in the columns and features in the rows.

Factor 3 captured signatures present in healthy subjects receiving broad-spectrum antibiotics, with a predominance of the closely related Streptococcaceae family *(Streptococcus* and *Lactococcus),* Lactobacillales order *(Lactobacillus* and *Granulicatella)* and Actinomyceteles order *(Actinomyces* and *Rothia).* Interestingly, all these bacteria have been shown to possess mutualistic properties with *Candida* in oral and vaginal environments, potentially through the modification of biofilm formation (Richard & Sokol, 2019; Arzmi *et al*, 2015; Kim *et al*, 2017; Uppuluri *et al*, 2017). These observations indicate that similar fungal-bacterial interactions are potentially maintained within the gastrointestinal tract, warranting further elucidation.

Notably, the majority of viral sequences that were associated with Factors 1 and 3 consisted of bacteriophages that significantly correlated with the presence of the corresponding bacterial targets in the same factor (**Fig. EV6**). The expansion of aerobic bacterial species during critical illness and following antibiotics can therefore potentially facilitate the enrichment of their corresponding bacteriophages (Shkoporov & Hill, 2019; Knowles *et al*, 2016). Other notable viral interactions were the increase of the mycoviruses Chrysovirus and Partitivirus–which are capable of infecting fungi (Ghabrial *et al*, 2015)- in healthy subjects following antibiotic exposure. These findings suggest that transkingdom interactions are occurring beyond intestinal bacteria, further underscoring the complexity of relationships within the intestinal environment.

An important indicator of the influence of the bacterial microbiota on the fungal population in the gut is the dramatic increase in the fungal burden after antibiotic treatment (Richard & Sokol, 2019). This phenomenon can partly be explained by antibiotic-induced alterations in nutrient availability, yet a loss of the direct inhibitory effects of anaerobic bacteria towards fungal expansion has also been documented (Nguyen *et al*, 2011; Fan *et al*, 2015; García *etal,* 2017). Therefore, to further explore the association between functional and absolute profiles of the bacterial microbiome and fungal expansion, we performed targeted bacterial 16S rRNA and fungal 18S rRNA quantitative PCRs to calculate intestinal fungal/bacterial ratios, and simultaneously quantified the absolute abundance of the SCFAs butyrate, acetate and propionate using Nuclear Magnetic Resonance (NMR) spectroscopy. We observed a strong depletion of SCFAs in both critical illness and following antibiotic perturbation, with the latter having the most significant impact (**Fig. 4a,b**). Notably, both conditions were associated with increased fungal/bacterial ratios, increasing as much as 103-104 times. This decrease of SCFAs coincided with a gradient of depletion along the axis of Factors 1 and 3 (**Fig. 4c**). Finally, we observed that absolute faecal SCFA concentrations were inversely correlated with absolute fungal copies, with propionate levels displaying the strongest correlation (r=0.75; p <0.0001, **Fig. 4d**). These data suggest that fungal expansion not only occurs in the context of decreased absolute bacterial abundance, but is also dependent on altered functions of the remaining bacterial communities in the intestinal environment.

**Figure 4:**
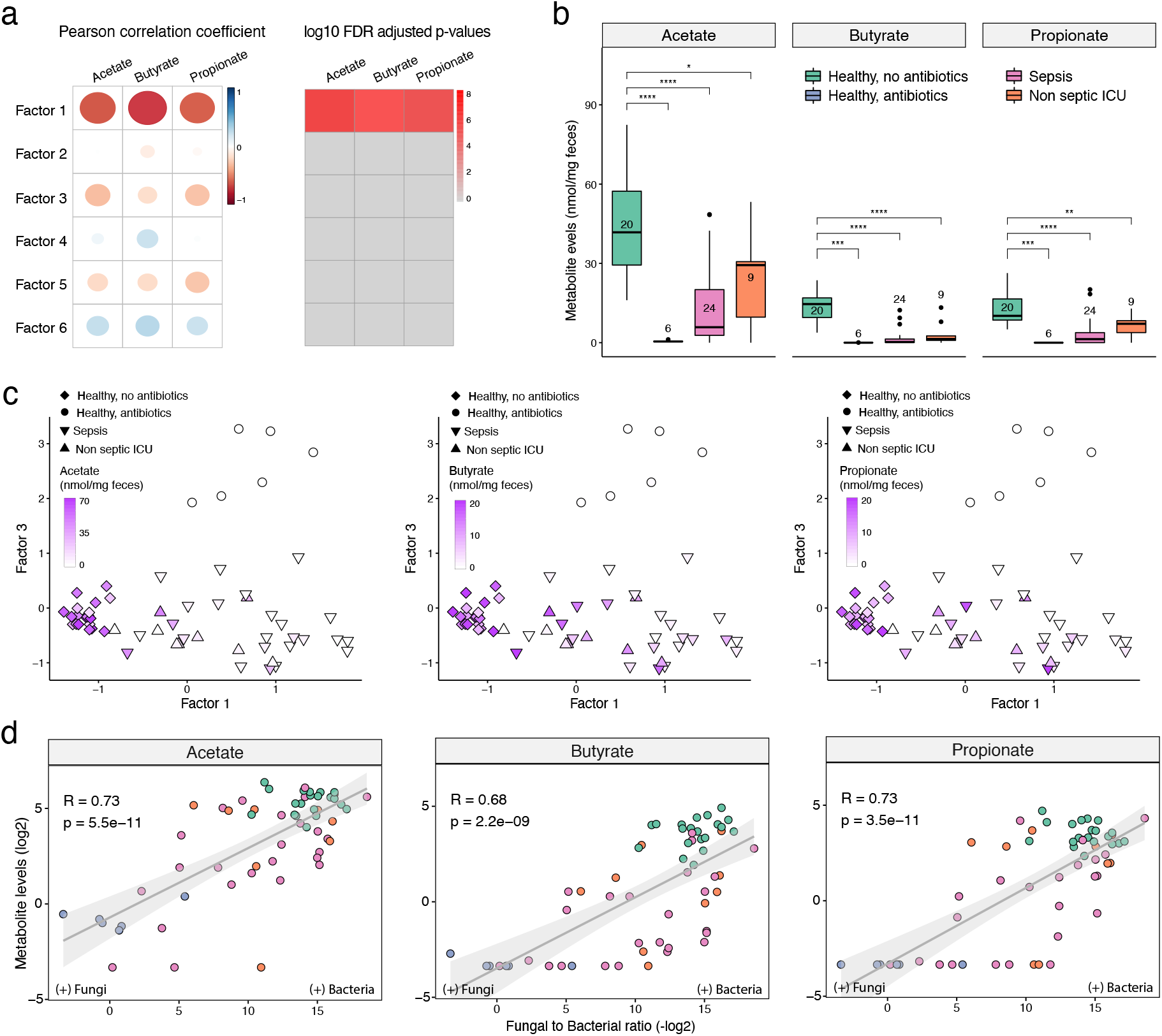
Correlation of total bacterial and fungal load with fecal levels of short-chain fatty acids in health and critical illness. (a) Association analysis between Factor values and SCFA levels. Left panel displays the Pearson correlation coefficient between factor values and the levels of three types of SCFA: butyrate, acetate and propionate. Right panel displays the corresponding FDR-adjusted and log-transformed p-values. (b) Box plots showing the SCFA concentrations (in mg per mg of feces, y-axis) per sample group (x-axis). In the box plots, the central rectangle spans the first quartile to the third quartile (the interquartile range or IQR), the central line inside the rectangle shows the median, and whiskers above and below the box. Given the non-parametric nature of the data, p values were calculated using the Wilcoxon rank sum test. (c) Scatter plot of Factor 1 (x-axis) versus Factor 3 (y-axis) values. Each dot represents a sample, shaped by the sample group and coloured by SCFA concentrations (in mg per mg of feces). (d) Scatter plot of Fungal-to-Bacterial absolute levels ratio (after log10 transformation, x-axis) versus SCFA concentrations (after log2 transformation, y-axis). The line represents the linear regression fit and the shade the corresponding 95% confidence interval. Corresponding Pearson correlation coefficients and p-values are also displayed in the top left corner.

In conclusion, our findings shed light into the dynamics and shared variations between kingdoms following broad-spectrum antibiotic modulation and critical illness. The short- and long-term impact of these disruptions will be an important focus of future investigations.

## Materials and Methods

### Study design and participants

Patients were recruited as part of a large prospective observational study in critically ill patients admitted to the ICU (Molecular Diagnosis and Risk Stratification of Sepsis (MARS) study; clinicaltrials.gov identifier NCT01905033) (van Vught *et al*, 2016; Lankelma *et al*, 2017b). A total of 33 randomly selected adult patients who were admitted to the ICU of the Academic Medical Centre (Amsterdam, The Netherlands) between October 2012 and November 2013 were included. Patients who were transferred from other ICUs or had an expected length of ICU stay of <24 h were excluded. All patients met at least two of the following criteria: body temperature of ≤36 or ≥38 °C, tachycardia of >90 /min, tachypnoea of >20 /min or partial pressure of carbon dioxide (pCO2) of < 4.3 kPA, and leukocyte count of <4 × 10E9/L or >12 × 10E9/L. Sepsis was defined when the inclusion criteria were associated with suspected infection within 24 hours after ICU admission, with subsequent systemic therapeutic administration of antibiotics to the patient (Lankelma *et al,* 2017b). The control group consisted of 13 healthy, non-smoking human subjects who had not taken antibiotics during the previous year (clinicaltrials.gov identifier NCT02127749) (Haak *et al*, 2019; Lankelma *et al*, 2017a). Six healthy subjects received oral broad-spectrum antibiotics (ciprofloxacin 500 mg q12h, vancomycin 500 mg q8h and metronidazole 500 mg q8h) for seven days. Subjects were asked to collect faecal samples before antibiotic treatment and one day after the 7-day course of antibiotics. Fresh stool samples by ICU patients were stored at 4 °C and transferred to −80 °C within 24 hours of collection. Faecal samples by healthy subjects were collected in plastic containers, stored at −20 °C at home and were transported to the study centre for storage at −80 °C within 24 hours. Written informed consent was obtained from all healthy subjects and patients or their legal representative. Ethical approval for both the patient and healthy subject studies was received from the Medical Ethics Committee of the Academic Medical Centre in Amsterdam, and all research was conducted in accordance with the declaration of Helsinki.

### Bacterial and fungal microbiota sequencing

Faecal DNA was extracted and purified using a combination of repeated bead-beating (method 5) (Costea *et al*, 2017) and the Maxwell 16 Tissue LEV Total RNA Purification Kit (Promega, Maddison, WI, USA), with STAR (Stool transport and recovery) buffer (Roche, Basel Switzerland). Negative extraction controls (DNA-free water) were processed in a similar manner.

Twenty nanograms of DNA was used for the amplification of the bacterial 16S rRNA gene with V3-V4 341F forward and 805R reverse for 25 cycles. The PCR was performed in a total volume of 30 μl containing 1× HF buffer (Thermo Fisher Scientific, Waltham, MA, USA), 1 μl dNTP Mix (10 mM; Promega, Leiden, the Netherlands), 1 U of Phusion Green High-Fidelity DNA Polymerase (Thermo Fisher Scientific, Waltham, MA, USA), 500 nM of the forward 8-nt sample-specific barcode primer containing the Illumina adapter, pad and link (341F (5’-CCTACGGGNGGCWGCAG-3’) 500 nM of reverse 8-nt sample-specific barcode primer containing the Illumina adapter, pad and link (805R (5’ GACTACHVGGGTATCTAATCC-3’)) 20 ng/μl of template DNA and nuclease free water. The amplification program was as follows: initial denaturation at 98 °C for 30 s; 25 cycles of denaturation at 98 °C for 10 s, annealing at 55 °C for 20 s, elongation at 72 °C for 90 s; and an extension at 72 °C for 10 min (Kozich *et al*, 2013). The size of the PCR products (~540 bp) was confirmed by gel electrophoresis using 4 μl of the amplification reaction mixture on a 1% (w/v) agarose gel containing ethidium bromide (AppliChem GmbH, Darmstadt (Germany).

Fungal composition was determined by ITS1 amplicon sequence analysis. PCR generated amplicon libraries were obtained from 100 ng faecal DNA using the ITS1 primer set containing an overhang for the Illumina Nextera platform [forward] 5’-TCGTCGGCAGCGTCAGATGTGTATAAGAGACAGCTTGGTCATTTAGAGGAAGTAA and [reverse] 5’-GTCTCGTGGGCTCGGAGATGTGTATAAGAGACAGGCTGCGTTCTTCATCGATGC primers and Phusion High Fidelity DNA Polymerase (Thermo Fisher Scientific, Waltham, MA, USA). A duplicate reaction in 20 μl was performed with following thermocycling conditions: initial denaturation at 98 °C for 1 min followed by 35 cycles denaturation (20 s), annealing (20 s at 58 °C) and extension (60 s at 72°C) and final extension at 72 °C for 5 min. The duplicates were pooled to a final volume of 40 μl. The PCR products were purified with AMPure XP beads (Beckman Coulter, Brea, CA, USA) and taken into 15 μl DNA-free water. A second amplification step was used to introduce multiplex indices and Illumina sequencing adapters using the Kapa polymerase system. A 24-cycle amplification reaction in 40 μl was performed with following conditions: initial denaturation at 95 °C for 3 min followed by denaturation (20s at 98 °C), annealing (20 s at 60 °C) and extension (60 s at 72 °C) and final extension at 72 °C for 5 min.

Bacterial and fungal PCR products were purified using AMPure XP beads (Beckman Coulter, Brea, CA, USA). Amplicon DNA concentration was measured with the Qubit fluorometric Quantitation method (Thermo Fisher Scientific, Waltham, MA, USA) and DNA quality was determined with the Agilent Bioanalyzer DNA-1000 chip, after which the purified products were equimolarly pooled. The libraries were sequenced using an Illumina MiSeq platform (GATCBiotech, Konstanz, Germany) using V3 chemistry with 2 × 251 cycles. Forward and reverse reads were truncated to 240 and 210 bases respectively and merged using USEARCH (Edgar, 2010). Merged reads that did not pass the Illumina chastity filter, had an expected error rate higher than 2, or were shorter than 380 bases were filtered. Amplified Sequence Variants (ASVs) were inferred for each sample individually with a minimum abundance of 4 reads (Callahan *et al*, 2016). Unfiltered reads were than mapped against the collective ASV set to determine the abundances. Bacterial taxonomy was assigned using the RDP classifier (Wang *et al*, 2007) and SILVA 16S ribosomal database V132 (Quast *et al*, 2013). Fungal taxonomy was assigned using the UNITE database (Nilsson *et al,* 2019).

### Viral microbiota sequencing and analysis

The collected faecal suspension was centrifuged to pellet cells and debris, and nucleic acids in the supernatant were extracted using the Boom method (Boom *et al*, 1990), followed by reverse transcription with non-ribosomal random hexamers (Endoh *et al*, 2005) and second strand synthesis. DNA was digested with Msel (T^TAA; New England Biolabs, Ipswich, MA, USA) and ligated to adapters containing a sample identifier sequence. Next, size selection with AMPure XP beads (Beckman Coulter, Brea, CA, USA) was performed to remove small DNA fragments prior to a 28-cycle PCR using adaptor-annealing primers. Small and large size selection was performed with AMPure XP beads to select DNA-strands with a length ranging between 150 and 550 nucleotides. Libraries were analysed using the Bioanalyzer (High Sensitivity Kit, Agilent Genomics, Santa Clara, CA, USA) and Qubit (dsDNA HS Assay Kit, Thermo Fisher Scientific, Waltham, MA, USA) instruments to quantify DNA length and concentration, respectively. Sample libraries were pooled at the equimolar concentration. In total, 50 pmol DNA of the pool was clonally amplified on beads using the Ion Chef System (Thermo Fisher Scientific, Waltham, MA, USA) and sequencing was performed on the Ion PGM™ System (Thermo Fisher Scientific, Waltham, MA, USA) with the ION 316 Chip (400 bp read length and 2 million sequences expected per run).

VIDISCA-NGS reads were aligned using BWA-MEM (Li, 2013) to a reference database consisting of the human reference genome (hg38), the SILVA SSU V132 database(Quast *et al*, 2013)qu, and all RefSeq viral genomes (downloaded in September 2019). Mapping outputs were further processed using the PathoID module of PathoScope 2.0 (Hong *et al*, 2014; Byrd *et al*, 2014), to reassign reads with multiple alignments to their most likely target. Viral candidates were aligned back to the reference database with BLASTn, and those aligning at ?95% for 100 bp were retained as hits. To ensure that all known eukaryotic viruses were detected with this approach, all reads that remained unmapped in the BWA-MEM step were analysed with a separate virus discovery bioinformatic pipeline, described in detail elsewhere (Kinsella *et al*, 2019). Briefly, rRNA reads were identified with SortMeRNA v2.1, non-rRNA reads were made non-redundant using CD-HIT v4.7, and these were queried against a eukaryotic virus protein database using the UBLAST algorithm provided as part of the USEARCH v10 software package (Edgar, 2010). Reads with a significant alignment to a viral protein were subsequently aligned to the non-redundant nucleotides database using BLASTn. Those with a best hit to a viral sequence were regarded as confidently viral, those not aligning to any sequences were regarded as putatively viral, while those with a non-viral best hit were regarded as false positives.

### Targeted measurement of intestinal protozoa

Automated nucleic acid extraction was performed on the MagNA Pure 96 instrument (Roche Applied Science, the Netherlands) according to the manufacturer’s protocol. DNA was eluted in 100 μl elution buffer (Roche Applied Science). Phocine Herpes Virus (PhoHV) DNA was added to all samples as an internal control for extraction and amplification efficiency. Presence of *Giardia lamblia, Cryptosporidium parvum, Entamoeba histolytica, Blastocystis hominis* and *Dientamoeba fragilis* was assessed by real-time PCR targeting the small subunit ribosomal RNA gene (SSU-rDNA) (van Hattem *et al*, 2019). Positive controls consisting of a plasmid containing the target sequence were included in every run, as well as negative extraction controls and negative PCR controls. Subjects were excluded from further analyses if internal controls tested negative in one or more samples.

### Targeted measurement of short-chain fatty acids

Sample preparation of faecal extracts and Nuclear Magnetic Resonance (NMR) spectroscopy for quantification of SCFAs was performed as described in Kim HK et al (Kim *et al*, 2018), with some modifications. Briefly, aqueous extracts of faeces were prepared by mixing 50-100 mg of faeces and 0.3 mL of deionized water, followed by mechanical homogenization in a Bullet Blender 24 (Next Avance Inc, Troy, NY, USA). The faecal slurry was centrifuged twice at 18213 × g for 10 min at 4 °C and 0.225 mL of the supernatant was mixed with 0.025 mL 1.5 M potassium phosphate buffer (pH 7.4) containing 2 mM sodium azide and 4 mM sodium trimethylsilyl-propionate-d4 (TSP-d4) in D2O. For each sample, one 1D 1H-NMR spectrum was acquired in a 14.1 T Avance II NMR (Bruker Biospin Ltd, Karlsruhe, Germany). Quantification of SCFAs from the NMR spectra was performed in ChenomX (Chenomx NMR suite 8.4) using the known concentration of TSP-d4.

### Quantitative PCR for bacterial and fungal load

For the measurement of total bacterial content in faecal samples, we used the method as reported by Nadkami and colleagues (Nadkarni *et al*, 2002), with modifications. Briefly, we used a primer concentration of 500 nM in a final volume of 10 μl with the SensiFast SYBR No-ROX Kit (Bioline, London, UK). The amplification conditions were as follows: initial denaturation at 95 °C for 5 s followed by denaturation (10 s at 95 °C) – annealing (10 s at 66 °C) – extension (20 s at 72 °C) for 44 repetitive cycles in a BioRad CFX96 thermocycler (Hercules, CA, USA). The primerset of FungiQuant (Liu *et al*, 2012) was used for fungal load determination, with modifications. The final PCR primer concentration was 500 nM in a volume of 10 μl with the SensiFast SYBR No-ROX Kit (Bioline, London, UK). The following amplification program was used: initial denaturation at 95 °C for 5 s min followed by denaturation (10 s at 95 °C) annealing (10 s at 60 °C) and extension (20 s at 72 °C) in 44 repetitive cycles in a BioRad CFX96 thermocycler (Hercules, CA, USA). Following amplification, fungal and bacterial ratios were calculated using LinRegPCR (Ruijter *et al*, 2009).

### Multi-Omics Factor Analysis (MOFA)

The input to MOFA is a list of matrices with matching samples, where each matrix represents a different data modality. Bacterial 16S rRNA ASVs, fungal ITS1 rRNA ASVs and viral sequences were defined as separate data modalities. As a filtering criterion, bacterial and fungal features were required to have a minimum of 10 ASVs observed in at least 25% of the dataset. In addition, to mitigate the sparsity of the data and to simplify the interpretation, we collapsed the inferred bacterial and fungal ASVs and viral sequences to their respective Family or Genus level. The number of sequences were subsequently scaled using a centralized-log-ratio (Aitchison, 1982), which has shown to be effective in normalizing compositional data (Gloor *et al*, 2017).

Model inference is performed using variational Bayesian inference with mean-field assumption (Argelaguet *et al*, 2018). The resulting optimisation problem consists of an objective function that maximises the data likelihoods (i.e. the variance explained) under some sparsity assumptions (Argelaguet *et al*, 2019b) which yields a more interpretable model output.

After model fitting, the number of factors was estimated by requiring a minimum of 5% variance explained across all microbiome modalities. The downstream characterization of the model output included several analyses:

- Variance decomposition: quantification of the fraction of variance explained (R2) by each factor in each view, using a coefficient of determination (Argelaguet *et al*, 2019b, 2018, 2019a).
- Visualization of weights: the model learns a weight for every feature in each factor, which can be interpreted as a measure of feature importance. Larger weights (in absolute value) indicate higher correlation with the corresponding factor values. The sign of the weight indicates the directionality of the variation: features with positive weights are positively associated with the corresponding values, whereas features with negative weights are negatively associated with the corresponding values.
- Visualization of factors: each MOFA factor captures a different dimension of heterogeneity in the microbiome composition. Mathematically, each factor ordinates cells along a one-dimensional axis centred at zero. Samples with different signs manifest opposite phenotypes along the inferred axis of variation, with higher absolute value indicating a stronger effect. Note that the interpretation of factors is analogous to the interpretation of the principal components in PCA.
- Data reconstruction: MOFA generates a compressed low-dimensional representation of the data. By taking the product of the factors and the weights, the model can reconstruct a normally-distributed denoised representation of the input data. This is particularly useful for the visualisation of sparse readouts.

### Statistics

All analyses were performed in the R statistical framework (Vienna Austria, version 3.6.1). To assess alpha diversity and richness, we calculated the Shannon Diversity Index and Observed Taxa Richness index with the phyloseq package17. Data were not normally distributed and are therefore presented as median and interquartile range (IQR), while data were analysed using a Wilcoxon rank-sum test. Associations between Factor values and covariates were analysed using linear regression by Pearson correlation coefficients. Statistical significance is called at 10% FDR.

## Data availability

Raw sequencing data (bacterial and fungal ASVs, VIDISCA-NGS sequencing reads) will be submitted to the European Nucleotide Archive (ENA; accession number PRJEB37289) prior to publication. All code used for analysis is available at https://github.com/bwhaak/MOFA_microbiome. Links to the processed data are included in the GitHub repository.

## Acknowledgements

The authors acknowledge Lonneke A. van Vught, Maryse A. Wiewel, Friso M. de Beer, Lieuwe D.J. Bos, Gerie J. Glas, Roosmarijn T.M. van Hooijdonk, Michaëla A.M. Huson, Laura R.A. Schouten, Marleen Straat, Esther Witteveen, and Luuk Wieske (Department of Intensive Care, Amsterdam UMC, Location Academic Medical Center, the Netherlands) for their participation in data collection. The authors would also like to thank Jorn Hartman, Michelle Klein, Martin Deijs, Maarten Jebbink, Patricia Broekhuizen – van Haaften and Ellen Wentink – Bonnema for their indispensable help in the laboratory work on the faecal samples. This work was supported by the Netherlands Organization for Scientific Research (NWO; Vidi Grant 91716475).

## Author contributions

BWH and WJW conceived the original study. RA performed the MOFA+ analysis. CMK and CMvdH designed and performed the viral sequencing and bioinformatics pipeline. SK and MG designed and performed the NMR analyses. Fungal profiling and sequencing was designed and performed by WdJ and TBMH. Protozoal analysis was overseen by TG. Microbiome sequencing and initial analysis was performed and facilitated by RFK, FH and WMdV. MS and TvdP oversaw sample collection on the ICU, JML oversaw sample collection of the healthy subjects. BWH, RFK and RA analysed the data, wrote the original manuscript, and prepared the final figures. CMvdH and WJW secured funding for this project. All authors have seen and approved the final version of the manuscript.

