## Supplementary Figures for "Intestinal transkingdom analysis on the impact of antibiotic perturbation in health and critical illness"

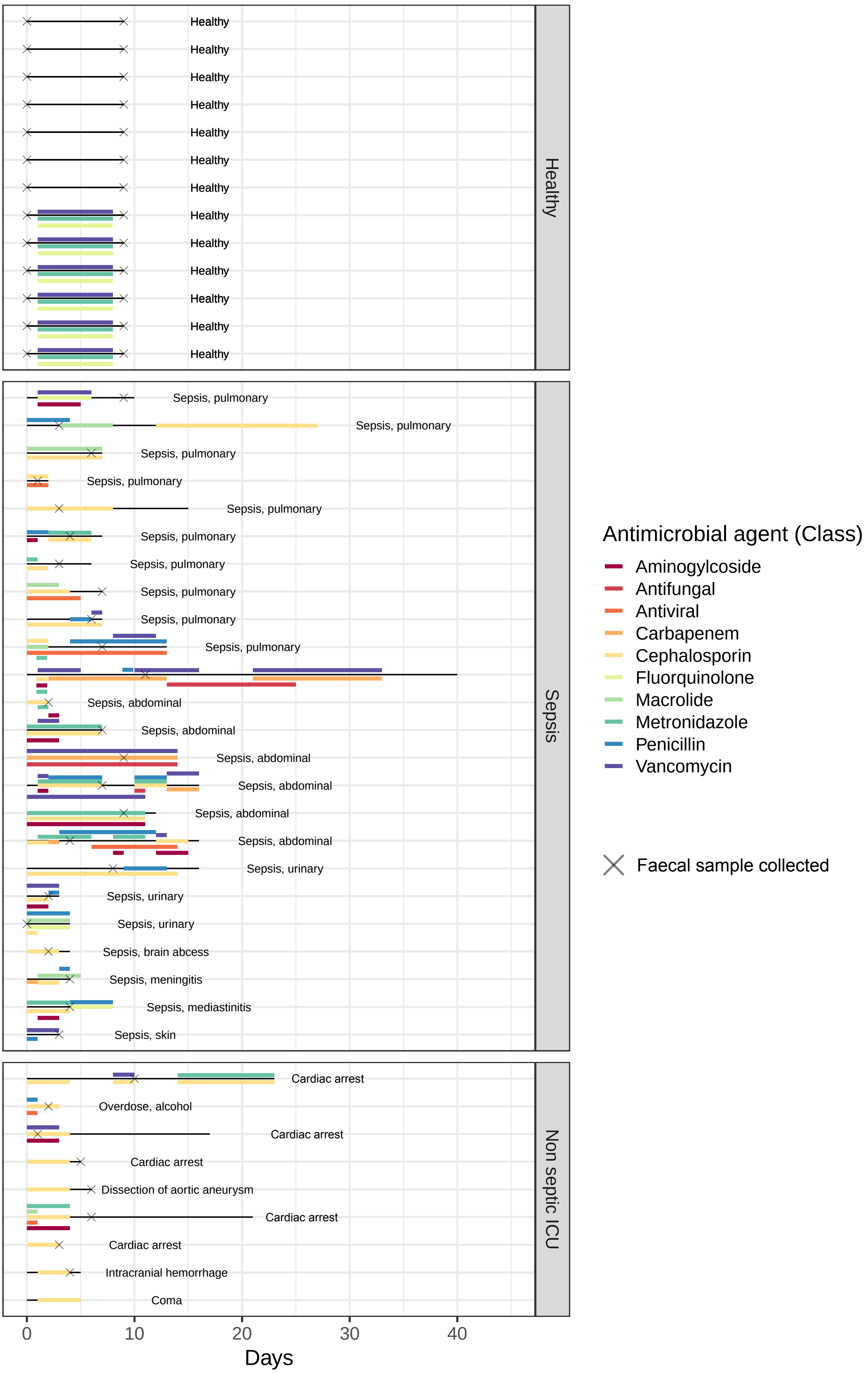


**Figure S1: Overview of antibiotic exposure of study cohort.**
Healthy volunteers are displayed at the top, septic patients in the middle and non-septic ICU patients are displayed at the bottom. Black line indicates the length of stay (for sepsis and non-septic ICU patients) or length of follow up for volunteers. Crosses indicate the days of faecal sample collection.

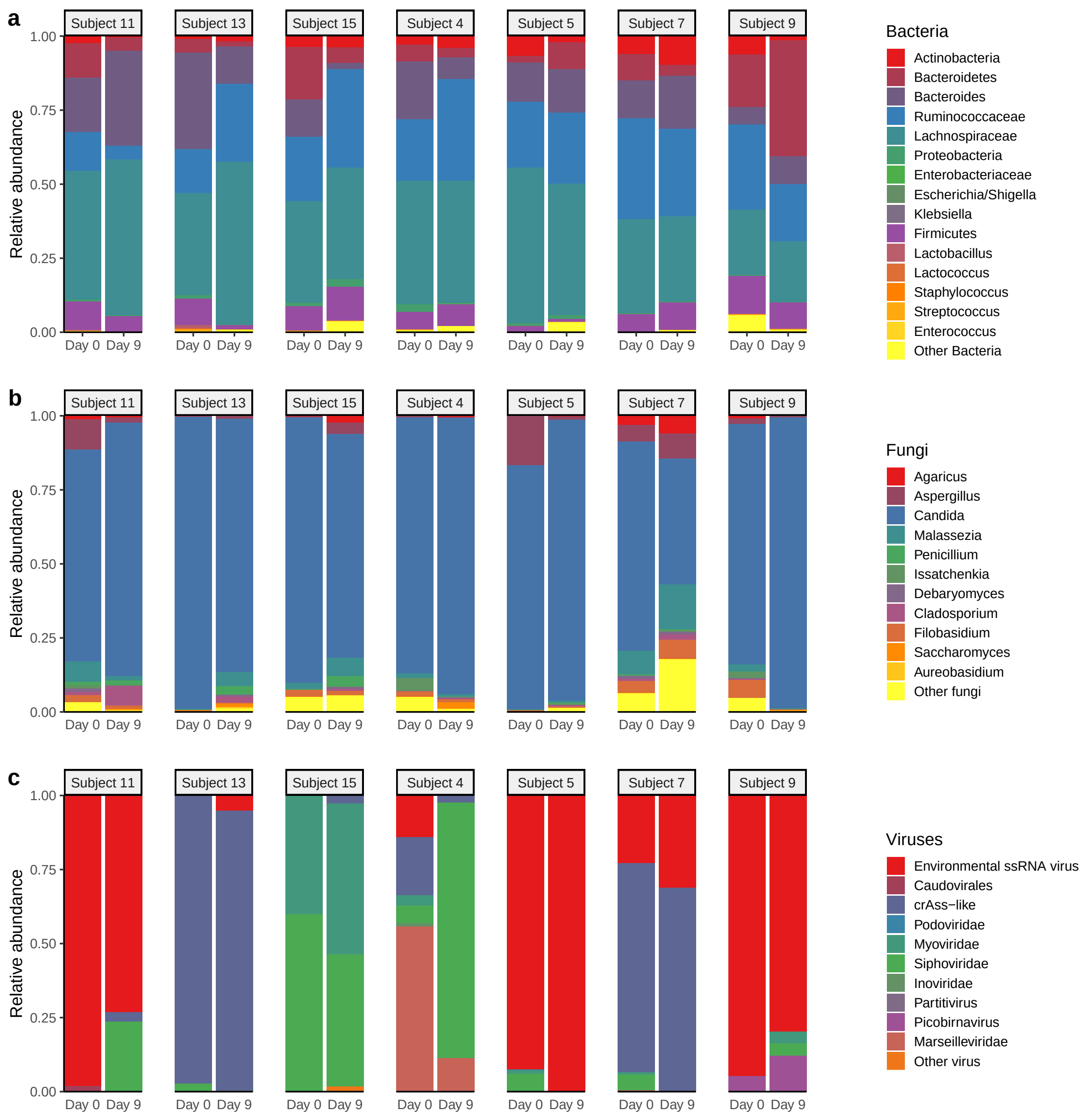


**Figure S2: Temporal stability of bacterial, fungal and viral microbiome In healthy volunteers.** Samples collected from healthy volunteers (n =7) not exposed to antibiotics on day 0 and day 9. Relative proportion of sequence reads at the Genus level assigned to different bacterial (a)and fungal (b) and viral (c) taxa.

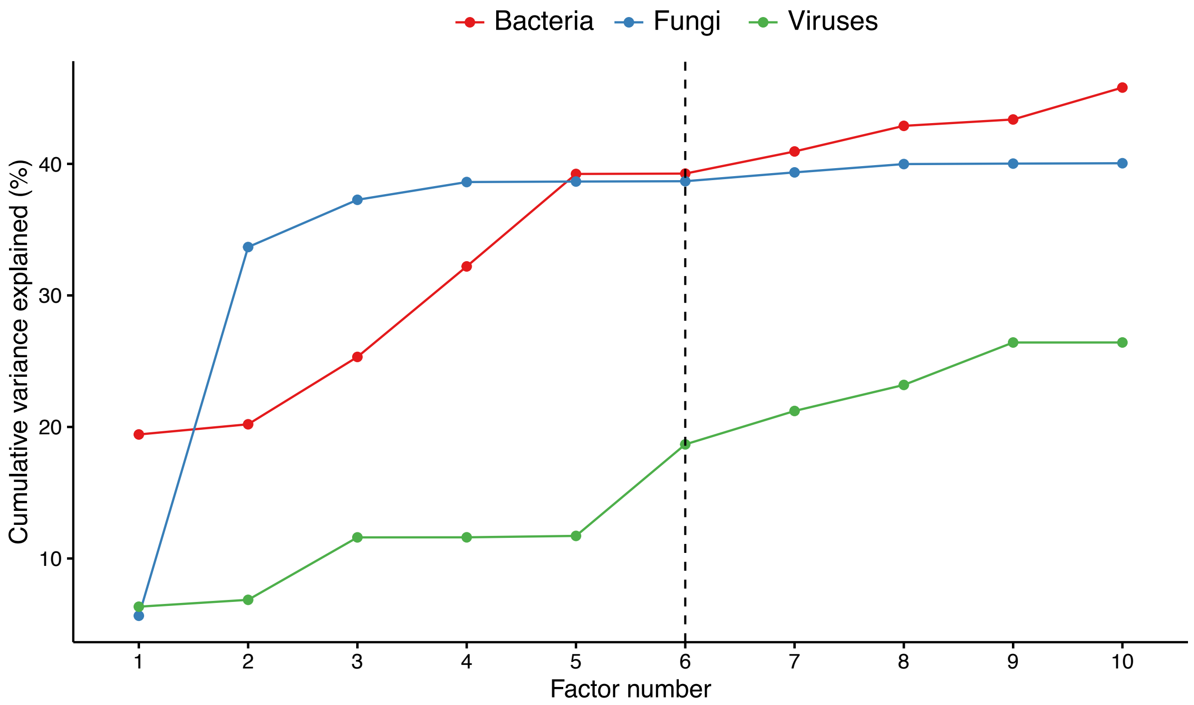


**Figure S3: Cumulative variance explained (per microbiome modality, y-axis) versus factor number (x-axis).**
The dashed line indicates the number of factors that were selected for downstream analysis (minimum of 5% variance explained across all data modalities).

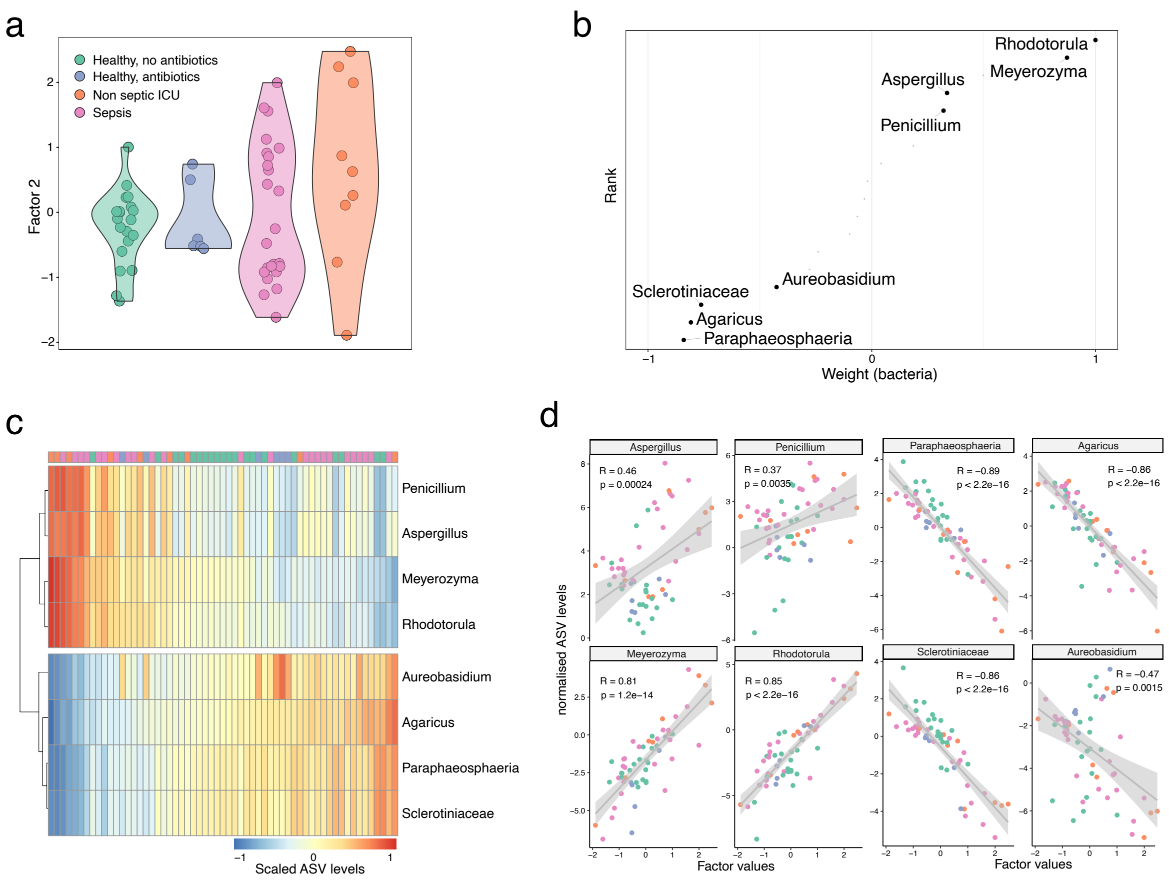


**Figure S4: Depiction of Factor 2, characterized by variation of low abundant fungi in critical illness.**

(a) Beeswarm and violin plots display Factor 2 values, indicating that the factor is driven by features that are predominantly present or absent during critical illness, with limited involvement within healthy volunteers.

(b) Distribution of Fungi weights for Factor 2. Labelled are representative fungi among the largest weights.

(c) Heatmap displaying the reconstructed and scaled ASV levels (see Methods) for the fungi labeled in (b).

(d) Scatter plot displays the association between factor values and normalised ASV levels. The line represents the linear regression fit and the shade the corresponding 95% confidence interval. Pearson correlation coefficients and p-values are displayed.

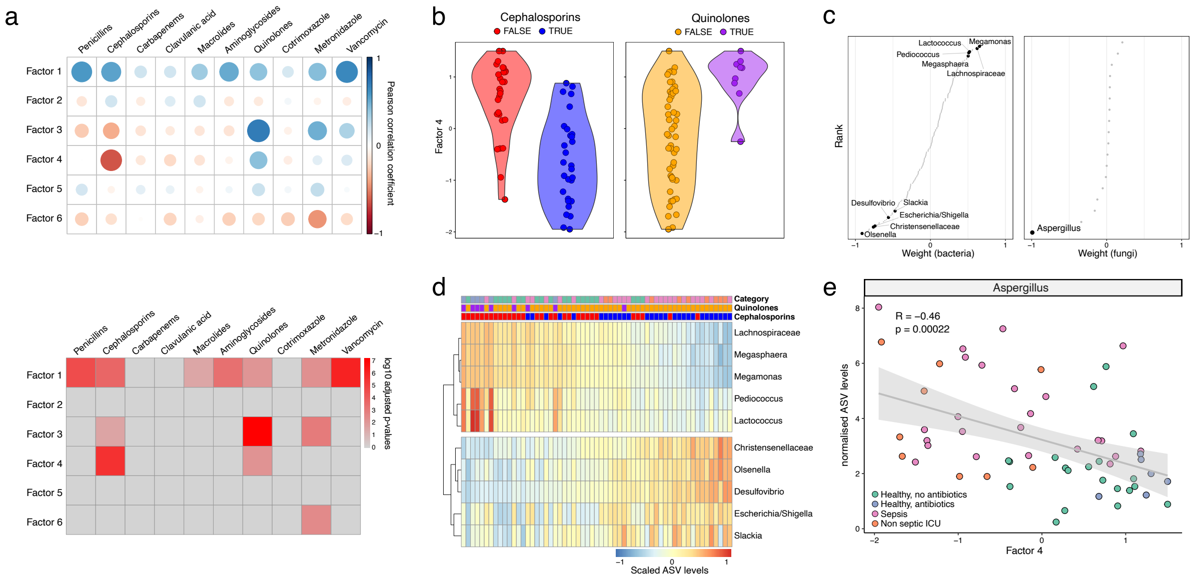


**Figure S5: Characterisation of Factor 4 as microbiome response to Cephalosporins and Quinolones.**

(a) Association analysis between Factors and antibiotic treatment. Top panel displays the Pearson correlation coefficient between factor values and the antibiotic treatment indicator variable. Bottom panel displays the associated log10 FDR-adjusted p-values.

(b) Beeswarm and violin plots display Factor 4 values, coloured by Cephalosporin (left) and Quinolone treatment (right).

(c) Distribution of Bacterial (left) and Fungi (right) weights for Factor 4. Labeled are representative bacteria and fungi among the largest weights.

(d) Heatmap displaying the reconstructed and scaled ASV levels (see Methods) for the bacteria labeled in (c).

(e) Scatter plot displays the association between factor values and normalised ASV levels in Aspergillus. The line represents the linear regression fit and the shade the corresponding 95% confidence interval. Pearson correlation coefficient and associated p-value is displayed in the top left corner.

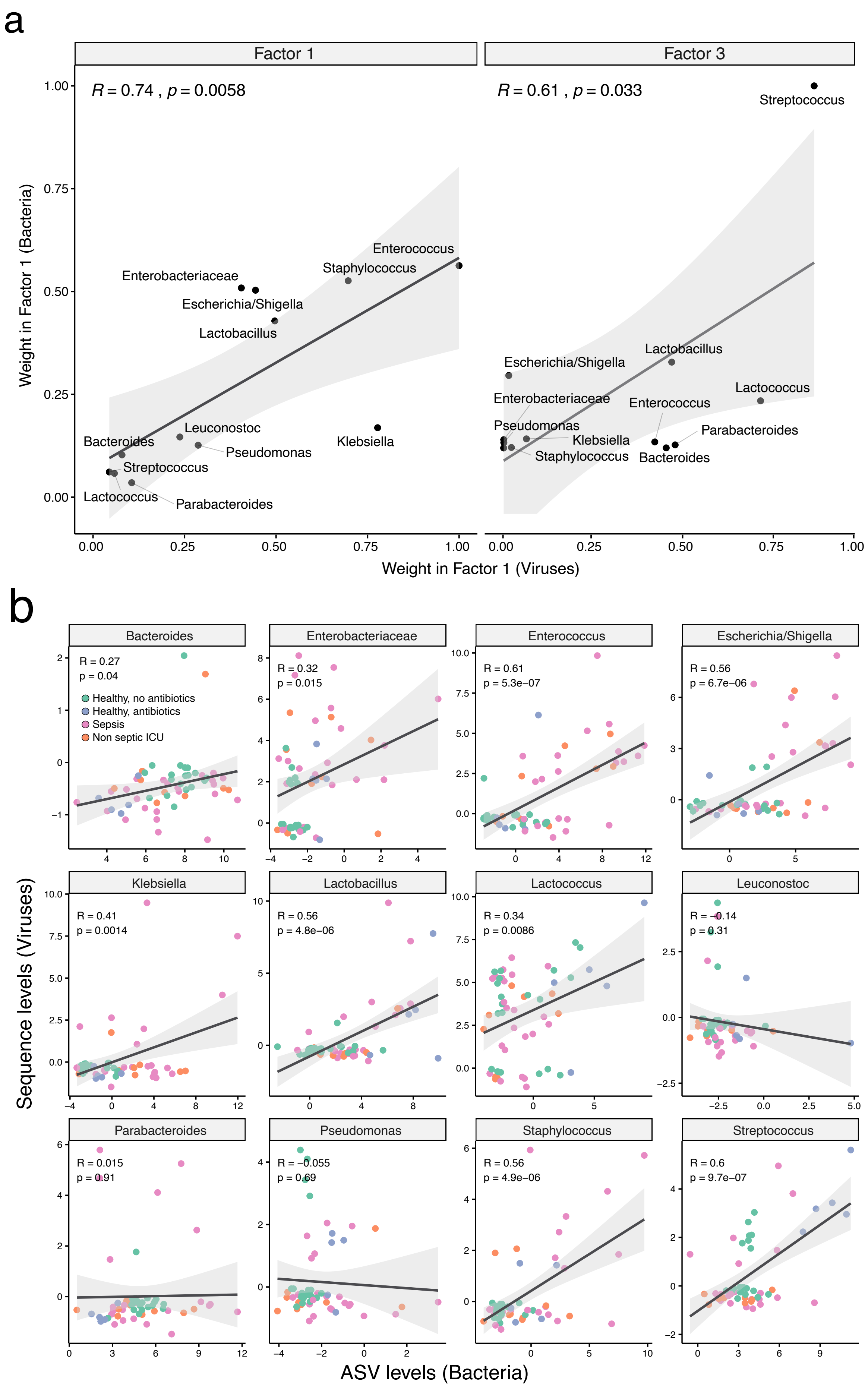


**Figure S6: Positive association between bacterial levels and associated viral phages.** (a) Scatter plots of the viral weights (x-axis) versus bacterial weights (y-axis), for MOFA Factor 1 (left) and Factor 3 (right). Each dot corresponds to a bacteria and its associated phage. (b) Scatter plots of the concentration levels of bacteria (x-axis) and the corresponding viral phages [illegible]
